# Parallel CLE peptide signaling pathways control nodulation in pea

**DOI:** 10.1101/2025.11.17.688984

**Authors:** Tiana E. Scott, Kate E. Wulf, Alejandro Correa-Lozano, Karen Velandia, James B. Reid, Eloise Foo

## Abstract

Legume root nodulation with nitrogen-fixing bacteria requires precise control via root-shoot-root autoregulation of nodulation (AON). Post-translationally modified root-derived CLAVATA3/Embryo Surrounding Region-Related (CLE) peptides signal through shoot acting leucine-rich repeat receptors (CLAVATAs) to regulate nodule number and this pathway is a target to optimise nodulation. We characterise the AON system in the crop model pea (*Pisum sativum* L.) and address key gaps in our understanding of AON; the role of parallel signalling pathways, shoot receptor complexes and downstream targets. We use novel mutant combinations, overexpression, grafting, gene expression and careful analysis of infection and nodule organogenesis using GFP-labelled rhizobium. These studies provide evidence that in pea both *Ps*CLE12 and *Ps*CLE13 require arabinosylation via *Ps*RDN1. Perception of *Ps*CLE12 and *Ps*CLE13 in the shoot to suppress the mature nodules in the root requires the pea CLAVATA1 orthologue *Ps*NARK and *Ps*CLV2. However, we found little evidence that *Ps*CLE12 and *Ps*CLE13 suppress infection thread development or that they act via *Ps*TML1 and/or *Ps*TML2, root acting suppressors of nodulation, indicating a role for additional CLE signals. Grafting and double mutant studies indicate that *Ps*NARK can act together with *Ps*CLV2, but also independently, to influences nodulation providing *in planta* evidence for shoot receptor complexes that control AON.

**Highlight:** We characterise the specific CLE peptide signalling pathway that the model crop legume pea uses to control the number of nitrogen-fixing nodules formed on the root.

## Introduction

Leguminous plants in the Fabid clade have evolved the ability to associate with nitrogen fixing bacteria, called rhizobia, in part by co-opting elements from the more ancestral associations with phosphorous-acquiring arbuscular mycorrhizal (AM) fungi (reviewed by Lebreton and Keller, 2024). One fascinating example of this cross over is the autoregulatory pathway, an internal feedback system that limits the amount of root colonised by the symbiont (reviewed by Roy and Muller, 2022; Scott *et al*., 2024). Elegant studies in legumes indicate plants limit colonisation by AM fungi and rhizobial bacteria via common signalling elements in a root-shoot-root signalling system (e.g. Catford, 2003). The autoregulation of mycorrhizae (AOM) pathway is also present in non-legumes to limit AM colonisation (Müller *et al*., 2019; Wang *et al*., 2021; Wulf *et al*., 2023), suggesting that it is a deeply conserved pathway. The model legume pea (*Pisum sativum* L.) has served as a key species since the 1980s for unravelling the physiology of the autoregulation of symbiosis pathways (e.g. Morandi *et al*., 2000; Postma *et al*., 1988), but our understanding of the molecular mechanisms that govern autoregulation of nodulation (AON) in pea is piecemeal.

Furthermore, fundamental questions remain across legumes regarding how regulatory elements of the AON system interact, in particular the role of various receptor complexes, the likelihood of parallel pathways and the precise stages of nodulation and downstream transcriptional targets that the AON system regulates. Given that the AON system is currently a target for crop improvement (e.g. Zhong *et al*., 2024), establishing the exact nature of AON in crop legumes is key. In this study, we characterised the key components of the AON system of the agriculturally significant legume crop *Pisum sativum* (pea).

Legumes activate the AON pathway to systemically regulate nodule number. The activation of cortical cell division during nodule formation stimulates the production of specific rhizobial-induced (Rh) CLAVATA3/ ENDOSPERM SURROUNDING REGION-related (CLE) peptides in the roots (e.g. Lebedeva *et al*., 2022; Li *et al*., 2009; Yoro *et al*., 2019b). The activity of some of these Rh-CLE peptides is enhanced following post-translational arabinosylation via hydroxyproline O-arabinosyltransferase (HPAT) enzymes that act in the root, including *Mt*RDN1 in *Medicago truncatula* (MtCLE12; Imin *et al*., 2018; Kassaw *et al*., 2017) and orthologue PLENTY in *Lotus japonicus* (Yoro *et al*., 2019a). The Rh-CLE peptides are perceived in the shoot and, at least in *Lotus*, exogenous CLEs have been shown to be transported via the xylem (Okamoto *et al*., 2013). Shoot acting leucine-rich repeat receptor-like kinases (LRR-RLKs), including CLAVATA1 (CLV1) orthologs (*Mt*SUNN, *Lj*HAR1, *Gm*NARK and *Ps*NARK/SYM29), *Lj*KLAVIER (*Lj*KLV), the pseudokinase CORYNE (CRN) and LRR-

Receptor-like protein (LRR-RLP) CLAVATA2 (CLV2), that lacks an intracellular kinase domain, are important regulators of nodule number (for review see Roy and Muller, 2022). Several of these LRR receptors have been shown to be required for perception of specific Rh-CLE peptides through application and/or overexpression studies (Hastwell *et al*., 2018; Imin *et al*., 2018; Kassaw *et al*., 2017; Okamoto *et al*., 2013). Numerous studies across legume species demonstrate that the disruption of orthologous HPAT enzymes and LRR receptors results in excessive nodule development (Kassaw *et al*., 2017; Nishimura *et al*., 2002; Searle *et al*., 2003). Furthermore, Rh-CLE peptides have been shown to suppress nodule number when overexpressed or ectopically applied, and can function interspecifically, as petiole feeding of soybean Rh-CLEs *Gm*RIC1a and *Gm*RIC2a could suppress nodule number in pea, and did so via pea LRR receptors *Ps*NARK and *Ps*CLV2 (Hastwell *et al*., 2019), suggesting that this system is highly conserved across species.

The perception of Rh-CLE peptides in the shoot leads to subsequent suppression of nodule number in the root via systemic signalling (e.g. Kosslak and Bohlool, 1984). Rhizobial inoculation downregulates microRNA miR2111 via CLV1 orthologs in both soybean and *Medicago* (Tsikou *et al*., 2018; Zhang *et al*., 2021). miR2111 acts as a systemic positive regulator of nodulation and may do so by downregulating *TML* (TOO MUCH LOVE) gene expression in the root. *TML1* and *TML2* encode F-box proteins that negatively regulate nodule number (Gautrat *et al*., 2019; Magori *et al*., 2009; Takahara *et al*., 2013). The transcription of *TML1* and *TML2* is upregulated during nodulation and this is influenced by Rh-CLE peptides (Chaulagain *et al*., 2023; Gautrat *et al*., 2019; Takahara *et al*., 2013). The precise connection between *TML1/2* transcription and the other AON signalling elements is unclear. In *Medicago*, Gautrat *et al*. (2019) reported that the Rh-CLE upregulation of *TML1/2* required the SUNN receptor but Schnabel *et al*. (2023) observed no change in *TML1/2* expression in *sunn-4* or *rdn1-2* mutant lines relative to wild type. However in *Lotus*, grafting Rh-CLE overexpression and double mutant studies with *har1* (*sunn* ortholog) and *tml1* did suggest these genes act in series to suppress nodulation (Nishida *et al*., 2016). In addition to these signals, in *Lotus* cytokinins produced in the shoot in response to CLE-HAR1 signalling may also act as a shoot-derived inhibitor of nodulation (Sasaki *et al*., 2014). Downstream targets of the AON pathway include Nod factor perception genes that may limit subsequent infection (Gautrat *et al*., 2019), and a feedback loop with key nodulation gene *NODULE INCEPTION* (*NIN*), which acts both up and downstream of AON (Soyano *et al*., 2014).

Despite AON emerging as a target for crop enhancement, clear gaps in our understanding of the AON system persist. Firstly, although a range of receptors have been implicated in AON, the evidence for specific receptor complexes that operate *in planta* to control nodule number is limited. Bimolecular fluorescence complementation assays in *Nicotiana benthamiana* leaves suggest *Mt*SUNN-*Mt*CLV2-*Mt*CRN, *Mt*CLV2-*Mt*CRN and *Lj*HAR1 and *Lj*KLV associate to form heteromeric complexes (Crook *et al*., 2016; Miyazawa *et al*., 2010), although these studies do not supply direct evidence for complex formation *in planta*. The lack of an additive nodule phenotype reported in *har1 clv2* double mutants may indicate these receptors act together, although no quantitative data was presented (Krusell *et al*., 2011) and the nodulation phenotype of other double mutants in AON receptors has not been reported. Secondly, there is mounting evidence that the AON system is not a linear feedback loop, but likely to consist of several parallel pathways. For example, in *Lotus* the additive nodule number in *Ljplenty har1* double mutant plants suggests these genes act in at least partially parallel pathways (Yoro *et al*., 2019a). Furthermore, in *Medicago* the *Mtsunn* and *Mtrdn1* mutants examined using split root studies retained some ability to suppress nodule number, indicating other systemic pathways that also regulate nodule number (Kassaw et al., 2015), possibly the recently described CEP-CRA2 system (e.g. Luo *et al*., 2021). Thus, additional studies that probe both questions of receptor complexes and parallel pathways, via grafting between receptor mutants and examination of the nodulation phenotypes of double mutants in elements of the AON system, would be informative. Finally, the downstream signalling elements of AON and the stage of nodulation that the AON system targets it is not entirely clear. Although there are reports of some mutants or transgenics with altered AON signalling influencing infection thread number (Imin *et al*., 2018; Magori *et al*., 2009; Nishida *et al*., 2020; Wopereis *et al*., 2000; Yoshida *et al*., 2010), the vast majority of the studies only report nodule number. Therefore it is not clear if the AON system broadly influences infection or only nodule number.

Pea is as an ideal model species to investigate these gaps in our understanding of the AON system since it has characterised mutants in several key AON elements (CLV1-ortholog *Ps*NARK, *Ps*CLV2 and *Ps*RDN1), as well as two CLE peptides (*Ps*CLE12 and *Ps*CLE13) with key roles in limiting nodule number, possibly via suppression of *Ps*TML1/2 expression (Lebedeva *et al*., 2022). Furthermore, as a key legume crop, information about the AON system of pea offers potential targets for optimising nodulation for agriculture. In this study we establish a clear model of how AON genes interact in pea to regulate nodulation using gene expression studies, hairy root transgenics, mutants, grafting and careful quantification of infection and nodule number. We also highlight some key differences to the models established for the other model legumes, *Medicago* and *Lotus*.

## Materials and Methods

### Plant material

Pea genotypes were *Psnark* (P88, formerly *sym29*, disrupted in *PsCLV1*), *Psclv2* (P64, formerly *sym28*) and *Psrdn1* (P79, formerly *nod3*) derived from the parental line ‘Frisson’ (Duc and Messager, 1989; Krusell *et al*., 2002; Krusell *et al*., 2011; Sagan, 1996; Schnabel *et al*., 2011). Double mutants between *Psnark, Psclv2* and *Psrdn1* were selected in the F2 generation as outlined by Scott *et al*. (2024).

### Gene expression studies

For RNAseq analysis, wild type pea plants were grown in sterile conditions for 7 d (McAdam *et al*., 2017). For the 0 dpi treatment, the top three longest lateral roots from three plants were pooled and this was repeated 3 times to result in 3 biological replicates. The remaining plants were inoculated with *Rhizobium leguminosarum* symbiovar *viciae* strain RLV248 and at 10 dpi the top three longest lateral roots from three plants were pooled, this was repeated 5 times to result in 5 biological replicates. To monitor the expression of *PsCLE12* and *PsCLE13* following inoculation, wild type plants were grown as described above. The infection zone (2 cm of tissue taken 1 cm back from root tip) of the root was harvested 0, 2, 4 and 6 dpi as well as the mature zone 6 dpi (the next 2cm of root, where nodules form in pea;Velandia *et al*., 2024), with tissue pooled from 2 plants per replicate per time point. To examine nitrogen response, wild type pea was grown under sterile conditions, modified Long Ashton nutrient solutions with 0, 1mM or 10mM KNO_3_ was applied once a week and, when plants were 4 weeks old, 2-3 whole secondary roots were harvested per plant with two plants pooled per replicate. For all samples, RNA was extracted using ISOLATE II RNA Mini Kit (Bioline, Cincinnati, OH, USA). cDNA was synthetised from 1 μg of RNA using SensiFAST cDNA Synthesis Kit (Bioline). RT-qPCR was carried out in a Rotor-gene Q2 PLEX (Qiagen), using SensiMix SYBR Master Mix (Bioline), in duplicate for each biological replicate. The transcription factor TFIIa and actin genes were used to calculate the relative expression (Velandia *et al*., 2024). Primers are listed in Suppl Table 1.

Total RNA was extracted from whole root tissue samples and processed for mRNA library preparation (poly-A enrichment) which in turn was sequenced on a NovaSeq X Plus platform (PE150) at the Ramaciotti Centre for Genomics (University of New South Wales, Sydney, Australia). Quality control assessment of the RNA-seq data was performed using the FastQC tool (Andrews, 2017). Based on this analysis, reads were subsequently curated using the

Trimomatic module (Bolger and Giorgi, 2014) and remaining sequences underwent pseudoalignment against the pea (*Pisum sativum*) Cameor transcriptome (Alves-Carvalho *et al*., 2015) via the Kallisto algorithm (Bray *et al*., 2016). The differential expression analysis was done using the DESeq2 package (Love *et al*., 2014) that leverages the negative binomial distribution for analysing normalised transcript reads. The pairwise comparisons were done for normalised paired-end read data present at 0 days post inoculation (dpi, as a control), versus 10 dpi time points. The selection of DEGs was based on the false discovery rate corrected p-values under 0.05.

### Plant transformation studies

*PsCLE12* and *PsCLE13* under the control of a CAMV35S promoter were inserted in the pCAMBIA_CR1 (pCR1) vector that contains the DsRED transformation marker (Sevin-Pujol et al., 2017) using the Golden Gate Assembly Kit (New England Biolabs, Ipswich, MA, USA). PCR fragments for these genes were generated from pea root cDNA using primers containing the BsaI recognition site and matching overhangs (Table S1), following the manufacturer’s instructions and transferred into ARqua1 *Agrobacterium rhizogenes.* pCR1::CAMV35S::GUS plasmid or pCR1::*CASP*::GUS was used as a control.

Pea hairy root transformation was carried out with wild type (Frisson), *Psnark*, *Psclv2* and *Psrdn1* according to Velandia *et al*. (2024) and upon transfer to 1L sterile pots (50:50 gravel:vermiculite) plants were inoculated with *Rhizobium leguminosarum* symbiovar *viciae* strain labelled with GFP (RLV248G). Plants were grown at 20°C/15°C (day/night) in natural daylight for approximately 4 weeks. Peas were harvested and roots were screened under a fluorescence microscope to sort into transformed (DsRED positive) and untransformed roots (DsRED negative) and stored in 4% paraformaldehyde. For RNA extraction, 5-6 complete DsRED positive (transformed root) samples per genotype and construct combination were harvested and gene expression studies were carried out as outlined above to examine if *PsCLE12* and *PsCLE13* genes were overexpressed in transformed roots. To identify and measure nodulation structures, roots were examined using a Zeiss Axioscope5 fluorescence microscope. The total number of infection threads (IT), developing nodules and mature nodules were counted per cm of root length (1 each of DRED positive and DsRED negative assessed per plant). To estimate the total number of mature nodules on transformed (DsRED positive) and untransformed (DsRED negative) roots, all visible nodules were counted on 4-6 transformed and untransformed roots per genotype and construct combination. Roots were weighed and mature nodule number was expressed per gram of fresh weight (FW) or dry weight (DW) root.

### Grafting and double mutant studies

Epicotyl-epicotyl I-grafts were performed between wild type (Frisson), *Psnark* and *Psclv2* using growth conditions and techniques outlined in Foo *et al*. (2016). Plants were inoculated on the day of grafting with *Rhizobium leguminosarum* bv. *viciae* (RLV248) and received a weekly dose of modified Long Ashton nutrient solution with no nitrogen (N) and 5 mM NaH_2_PO_4_. Roots were harvested 4-5 weeks after grafting, nodule number on whole root system was counted and root, shoot and nodules were separated and dried at 55°C to obtain the DW. Nodule number was expressed per gram root DW.

For double mutant studies, wild type (Frisson), *Psnark*, *Psclv2, Psrdn1, Psnark clv2, Psnark rdn1* and *Psclv2 rdn1* plants were grown under sterile conditions as described by McAdam *et al*. (2017) for 16 days and inoculated with *Rhizobium leguminosarum* bv. *viciae* containing a GFP reporter (RLV248G) and received a weekly dose of modified Long Ashton nutrient solution with no N and 5 mM NaH_2_PO_4_. Root system was harvested 6 weeks after planting; 2-3 mature secondary roots from the midsection of the root system were harvested per plant and stored in 1% paraformaldehyde for infection scoring. One root per plant was assessed for the total number of infection threads (IT), developing nodules and mature nodules were counted per cm of root length, from 5 plants per genotype.

## Results

### Analysis of pea gene expression, including *PsCLE* genes

To examine the expression of AON genes, including *PsCLE* genes in response to rhizobial inoculation and nodulation, RNAseq was performed on whole wild type pea roots 0 and 10 dpi (days post inoculation). The reciprocal best hit approach was used to identify orthologues for *PsCLE* genes in pea. Sequences of *CLE* from *M. truncatula* and *Lotus japonicus* Lj1.0v1 (Karlo *et al*., 2020; Müller *et al*., 2019) were used as queries to search for the closest orthologues within the pea ‘Cameor’ transcriptome (Alves-Carvalho *et al*., 2015; Hastwell *et al*., 2019). Identified pea *CLE* genes were then subjected to a reciprocal BLAST search against the *M. truncatula* Mt4.0v1 and/or *L. japonicus* genome (https://phytozome-next.jgi.doe.gov). We identified several *PsCLE* genes not previously identified (PsCam026215, PsCam034760, PsCam045910, PsCam049008, PsCam049236, PsCam051274, PsCam060128 and PsCam008296) taking the total number of *PsCLE* genes to 56. The expression of 13 *PsCLE* peptides were significantly up-regulated by inoculation, including *PsCLE12* (PsCam040153) and *PsCLE13* (PsCam040702) (Fig. 1A), as has been observed for these two genes previously (Samorodova *et al*., 2018). The expression of four *PsCLE* genes was significantly downregulated in pea roots by inoculation (Fig 1A. The expression of other elements of the AON pathway was also examined, and we found the expression of *PsTML1* and *PsTML2* was significantly up-regulated 10 dpi (Suppl Table 2). Other genes that play important roles in nodulation, including Nod factor signalling, infection, nodule organogenesis and function are strongly upregulated by 10 days of exposure to rhizobia (Suppl Table 2).

**Figure 1.**
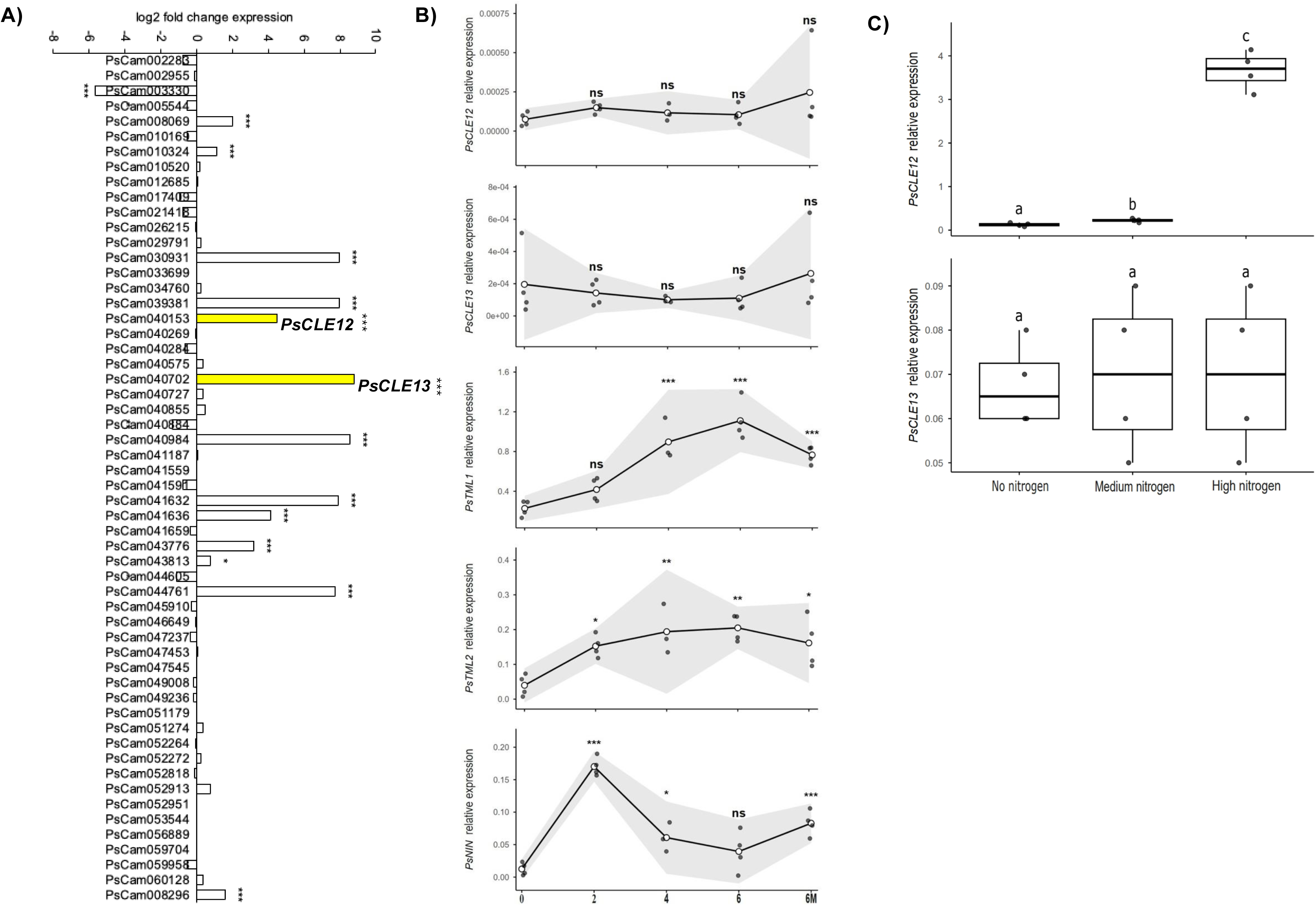
Expression profiles of key autoregulation genes in pea. (A) Log2-fold change in expression of *PsCLE* genes in wild type roots 10 days post inoculation (dpi) compared to 0 dpi, *PsCLE12* and *PsCLE13* are highlighted in yellow. Significant is stated * p-adj < 0.05,** p-adj <0.01, *** p-adj < 0.001 correspond to the p-value adjusted for false discovery ratio, n = 3-5. (B) Relative expression of *PsCLE12*, *PsCLE13, PsTML1, PsTML2 and PsNIN* in the infection zone of the root 0, 2, 4 and 6 dpi and in adjacent zone of roots forming nodules 6 dpi (6M). (C) Relative expression of *PsCLE12* and *PsCLE13* in whole secondary roots supplied weekly for 4 weeks with fertliser containing no nitrogen (no N), 1mM (medium N) or 10mM KNO_3_ (high N). For B and C, one-way ANOVA was performed and assumptions were tested by Shapiro–Wilk and Levene’s test. For B, the lines represent the mean ± 95% CI shown in grey. Dunnett’s test was used to compare each time point with time point zero. If assumptions were not met, Dunnett inference used HC3 standard errors. Significance is indicated as: ns p ≥ 0.05, * p < 0.05,** p<0.01, *** p < 0.001. For C, Tukey’s HSD post-hoc analyses was used, if assumptions were not met, a Welch ANOVA followed by the Games–Howell post hoc test was used. Letters indicate significant differences (p < 0.05).

The expression of *PsCLE12, PsCLE13, PsTML1, PsTML2* and *PsNIN* was monitored in roots in the days following inoculation and in response to nitrogen treatment (in the absence of rhizobium) by real time qPCR (Fig. 1B-C). No significant increase in *PsCLE12* and *PsCLE13* expression was observed in the infection zone of the root 2-6 dpi, although a small but not significant increase in expression was observed in the zone of the pea root where nodules become visible by 6 dpi (6M, Fig 1B,C; Velandia *et al*., 2024). Under our conditions, the expression of *PsCLE12* and *PsCLE13* only increased significantly at 10 dpi (Fig 1A). In contrast, *PsTML1* and *PsTML2* expression was significantly upregulated by 2 or 4 dpi relative to 0 dpi, with expression continuing to increase in the root infection zone until 6 dpi and also in the zone of the pea root where nodules become visible (6M; Fig. 1B). *PsNIN* was significantly upregulated at 2 – 4 dpi in the infection zone and was upregulated in the nodulation zone 6 dpi (6M; Fig. 1B). The expression of *PsCLE12* was significantly higher in plants treated with high nitrogen compared to medium or no nitrogen (Fig 1C), consistent with previous reports in pea examining short term response to nitrogen treatment (Lebedeva *et al*., 2022). However, in contrast to previous studies (Lebedeva *et al*., 2022), we did not find an increase in expression of *PsCLE13* in response to several weeks of high nitrogen (Fig 1C).

### *Ps*CLE12 and *Ps*CLE13 systemically suppress nodule number but not infection and act via *Ps*NARK and *Ps*CLV2

Previous studies have shown that overexpression of *PsCLE12* and *PsCLE13* in pea roots acts locally to suppress mature nodule number (Lebedeva *et al*., 2022). However, it was not clear if *PsCLE12* and *PsCLE13* acted systemically, if overexpression also influenced infection thread number and if these peptides act via putative receptors *Ps*NARK and *Ps*CLV2. To examine this, *PsCLE12* or *PsCLE13* were overexpressed (OE) in wild type, *Psclv2* and *Psnark* mutant plants (Fig. 2, 3). Within a genotype, both wild type and mutant roots transformed with *PsCLE12* overexpression vector had higher expression of *PsCLE12* compared to roots transformed with control construct (Suppl Fig 1A, B). A significant increase in expression of *PsCLE13* was also observed in transformed roots of *Psclv2* and wild type roots lines transformed with overexpression *PsCLE13* vector (Suppl Fig 2 B). High variability in *PsCLE13* expression occurred in the experiment with *Psnark* and wild type roots overexpressing *PsCLE13*, although a trend of increased expression in *PsCLE13* was observed in this experiment (Suppl Fig 2 A). Transformation did not appear to influence nodule number *per se* as, in plants transformed with control constructs (pCR1::CAMV35S::GUS or pCR1::CASP::GUS), both transformed and untransformed roots of the same plant had the same number of nodules (Fig 2C, Fig3C).

**Figure 2.**
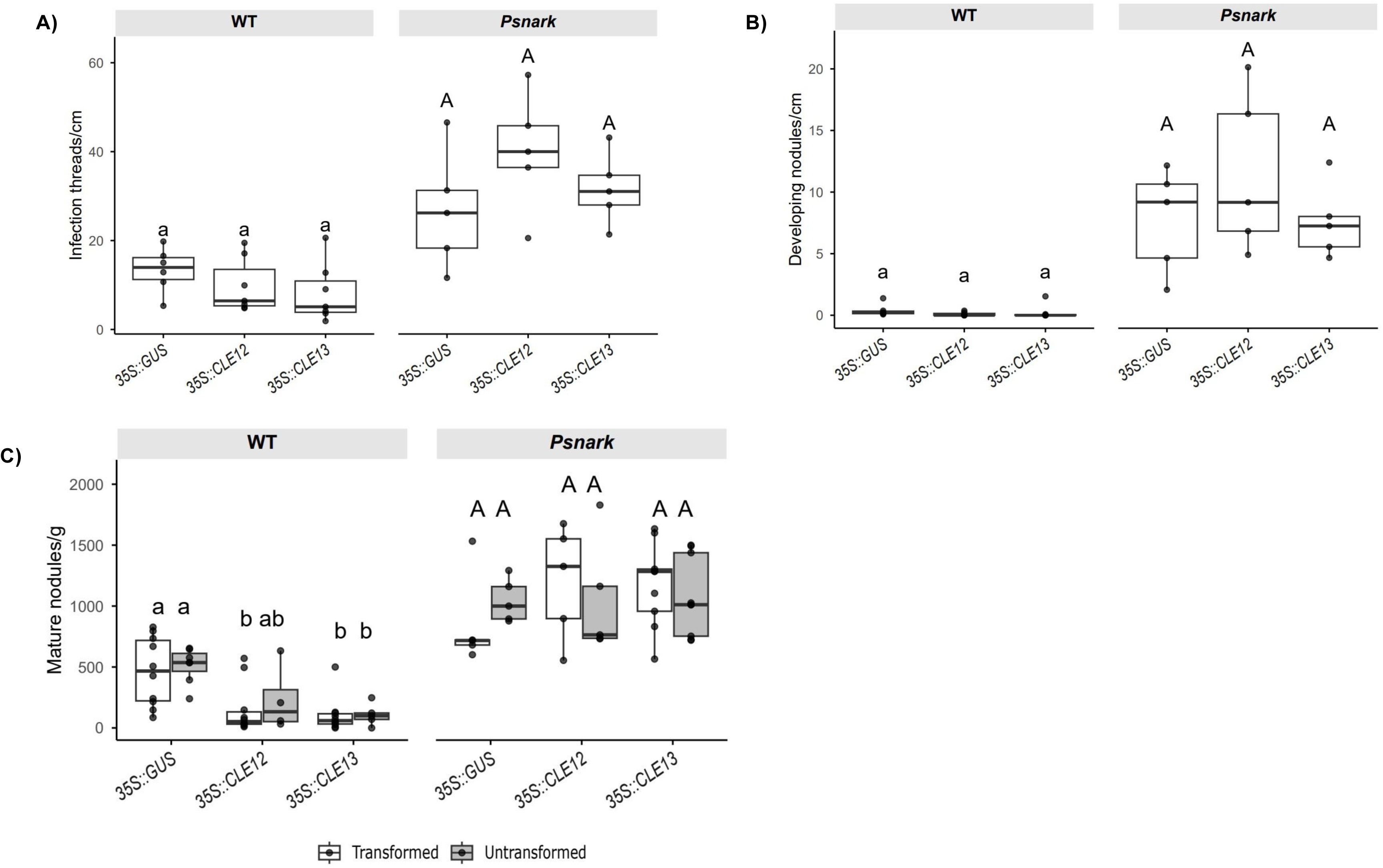
Overexpression of *Ps*CLE12 and *Ps*CLE13 suppresses nodulation in a *P*sNARK-dependant manner. Nodulation phenotype of wild type (WT; Frisson) and *Psnar*k mutant transformed via hairy root transformation with control (*35S::GUS*), *35S::PsCLE12* or *35S::PsCLE13* construct. (A) Number of infection threads per cm of transformed root, (B) number of developing nodules per cm of transformed root and (C) number of mature nodules per g fresh weight (FW) of transformed or untransformed root. Boxes show interquartile range, with the inner line representing the median of each treatment. Outliers fall beyond the whiskers, which encompasses up to 1.5 times the length of the interquartile range. Letters above each box represent significance values generated by one-way ANOVA and Tukey’s HSD post-hoc analyses comparing treatments within genotypes. (n= 5-10).

As observed in prior experiments without transformation (e.g. Foo *et al*., 2014), both *Psnark* and *Psclv2* controls displayed more nodules than wild type controls (Fig. 2C, Fig. 3C), and these mutants also displayed more infection threads and developing nodules (nodules only visible using microscope) than wild type control roots (Fig 2 A,B and Fig 3A,B). In both experiments, overexpression of *PsCLE12* or *PsCLE13* in wild type suppressed mature nodule number in both transformed and untransformed roots compared to wild type control roots (Fig 2C, Fig 3C). This indicates that *PsCLE12* and *PsCLE13* act systemically to suppress nodule number. Suppression of mature nodule number by *PsCLE12* or *PsCLE13* overexpression was not observed in *Psnark* or *Psclv2* mutant roots, whether transformed or untransformed, indicating these receptors are required for the systemic effect of *PsCLE12* and *PsCLE13* overexpression.

**Figure 3.**
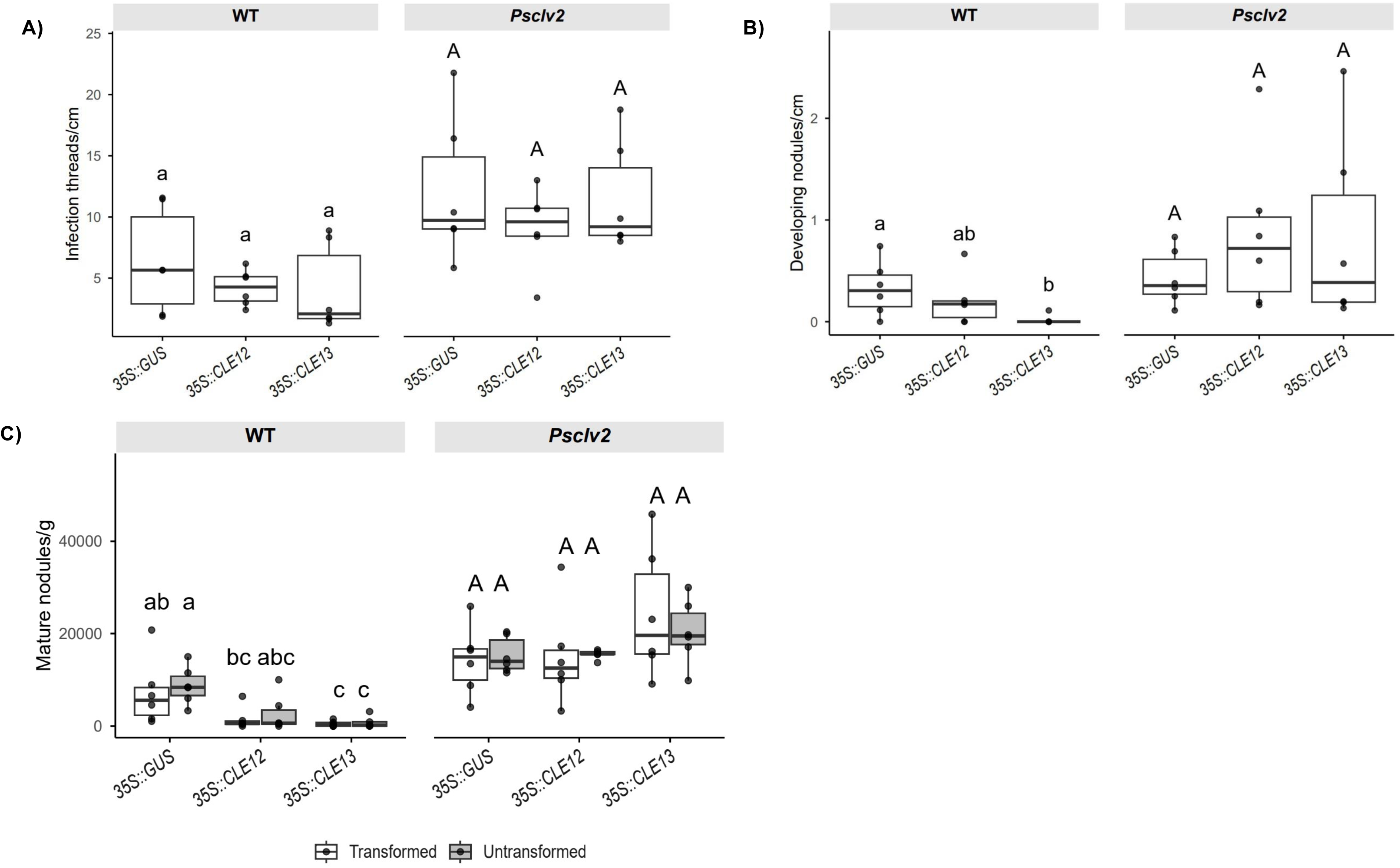
Overexpression of *Ps*CLE12 and *Ps*CLE13 suppresses nodulation in a *P*sCLV2-dependant manner. Nodulation phenotype of wild type (WT; Frisson) and *Psclv2* mutant transformed via hairy root transformation with with control (*35S::GUS*), *35S::PsCLE12* or *35S::PsCLE13* construct. (A) Number of infection threads per cm of transformed root, (B) number of developing nodules per cm of transformed root and (C) number of mature nodules per g dry weight (DW) of transformed or untransformed root. Boxes show interquartile range, with the inner line representing the median of each treatment. Outliers fall beyond the whiskers, which encompasses up to 1.5 times the length of the interquartile range. Letters above each box represent significance values generated by one-way ANOVA and Tukey’s HSD post-hoc analyses comparing treatments within genotypes. (n= 6).

The number of infection threads and developing nodules was also monitored in transformed roots. There was no significant effect of overexpression of *PsCLE12* or *PsCLE13* in any genotype on the number of infection threads formed in transformed roots compared to control transformed roots (Fig 2A, Fig 3A). In the experiment shown in Figure 2, very few developing nodules were observed in wild type plants, and no effect of overexpression of *PsCLE12* or *PsCLE13* observed (Fig 2B). In the experiment presented in Figure 3, there was a significant reduction in developing nodules in transformed wild type roots overexpressing *PsCLE13* compared to wild type control transformed roots (Fig 3B). As observed for mature nodule number, neither *Psnark* or *Psclv2* mutant roots overexpressing *PsCLE12* or *PsCLE13* displayed an altered number of developing nodules compared to control transformed roots (Fig2B, Fig 3B). Therefore, the overall lack of response of *Psnark* or *Psclv2* mutant roots overexpressing *PsCLE12* or *PsCLE13* suggests that *Ps*NARK and *Ps*CLV2 are receptors for *Ps*CLE12 and *Ps*CLE13.

Contrary to previous findings in pea (Lebedeva *et al*., 2022), we observed no increase in *PsTML1* or *PsTML2* expression in wild type overexpressing *PsCLE12* or *PsCLE13* relative to GUS control (Suppl Fig. 1, Suppl Fig. 2). Across two experiments, *PsTML1* and *PsTML2* expression was unchanged in *Psclv2* mutants compared to wild type but we did find an approximately 2-fold reduction in *PsTML1* and *PsTML2* expression in *Psnark* mutants (control and OE constructs) compared to wild type control (Suppl Fig. 1 A,B, Suppl Fig. 2 A,B). *PsNIN* expression was elevated in *Psnark* and *Psclv2* mutants (control and OE constructs) compared to wild type control. Overexpressing *PsCLE12* and *PsCLE13* significantly reduced *PsNIN* expression in wild type lines relative to *GUS* control in only one out of three experiments (Suppl Fig. 1, Suppl Fig 2).

### *Ps*CLE12 and *Ps*CLE13 likely require arabinosylation via *Ps*RDN1

Several rhizobial-induced CLE peptides have been shown to require arabinosylation to be active (Imin *et al*., 2018), and in some cases this is via modification by a HPAT enzyme encoded by RDN1 (Kassaw *et al*., 2017). To evaluate if *Ps*RDN1 is required for *Ps*CLE12 and/or *Ps*CLE13 to suppress nodule number, *PsCLE12* or *PsCLE13* was overexpressed in wild type and *Psrdn1* mutant plants (Fig. 4). Given the clear results presented in Fig 2 and Fig 3, infection thread number and untransformed root data was not reported in this experiment.

**Figure 4.**
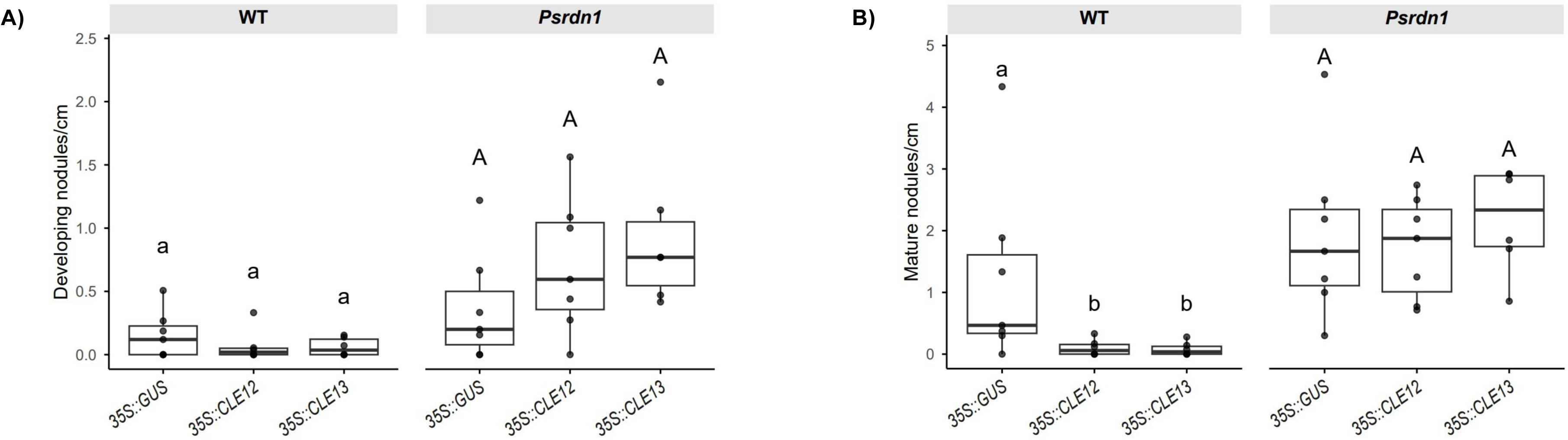
Overexpression of *Ps*CLE12 and *Ps*CLE13 suppresses nodulation in a *P*sRDN1-dependant manner. Nodulation phenotype of wild type (WT; Frisson) and *Psrdn1* mutant transformed via hairy root transformation with with control (*35S::GUS*), *35S::PsCLE12* or *35S::PsCLE13* construct. (A) Number of developing nodules per cm of transformed root and (B) number of mature nodules per cm of transformed root. Boxes show interquartile range, with the inner line representing the median of each treatment. Outliers fall beyond the whiskers, which encompasses up to 1.5 times the length of the interquartile range. Letters above each box represent significance values generated by one-way ANOVA and Tukey’s HSD post-hoc analyses comparing treatments within genotypes. (n= 6-7).

Within a genotype, both wild type and mutant roots transformed with overexpression vectors displayed significantly higher expression of *PsCLE12* or *PsCLE13* respectively (Suppl Fig. 1C, Suppl Fig. 2C). As expected, *Psrdn1* mutant roots exhibited higher numbers of developing and mature nodules compared to wild type roots (Fig. 4). A significant reduction in mature nodule number was recorded in wild type roots overexpressing *PsCLE12* or *PsCLE13* compared to wild type control roots (Fig. 4). This suppression was not seen in *Psrdn1* mutant plants (Fig. 4), suggesting *Ps*CLE12 and *Ps*CLE13 require arabinosylation by *Ps*RDN1 to be active.

Similar to *Psnark* and *Psclv2* experiments, *PsTML1* and *PsTML2* expression was not altered by *PsCLE12* or *PsCLE13* overexpression in wild type or *Psrdn1* mutant roots (Suppl Fig. 1C, Suppl Fig. 2C). Additionally, there was no apparent difference in *PsTML1* and *PsTML2* expression in *Psrdn1* mutants compared to wild type (Suppl Fig. 1c, Suppl Fig. 2c). In addition, *PsNIN* expression was not altered by either *PsCLE12* or *PsCLE13* overexpression or *Psrdn1* mutation relative to wild type (Suppl Fig. 1C, Suppl Fig. 2C).

### *Ps*NARK acts partly independently of *Ps*CLV2 to suppress nodulation, while *Ps*RDN1 likely acts in same pathway as *Ps*NARK and *Ps*CLV2

To examine if *Ps*NARK, *Ps*CLV2 and *Ps*RDN1 supress nodulation by acting in the same or parallel pathways, the nodulation phenotype of double mutant plants (*Psnark clv2, Psnark rdn1* and *Psclv2 rdn1*) was compared to wild type, *Psnark*, *Psclv2* and *Psrdn1* single mutant plants (Fig 5). In this experiment, *Psnark, Psclv2* and *Psrdn1* had only small but not significant increases in infection thread number compared to wild type (Fig 5 A). Double mutant *Psnark clv2* displayed significantly more infection threads than either *Psnark* or *Psclv2* single mutant plants (Fig 5 A, B?). This additive effect of disruption of both *PsNARK* and *PsCLV2* genes suggests these receptors act at least partially independently to suppress infection. In contrast, no additive effect on infection thread number was observed in *Psnark rdn1* or *Psclv2 rdn1* (Fig 5 A), indicating *Ps*RDN1 acts in same pathway as *Ps*NARK and *Ps*CLV2 to suppress infection thread formation.

**Figure 5.**
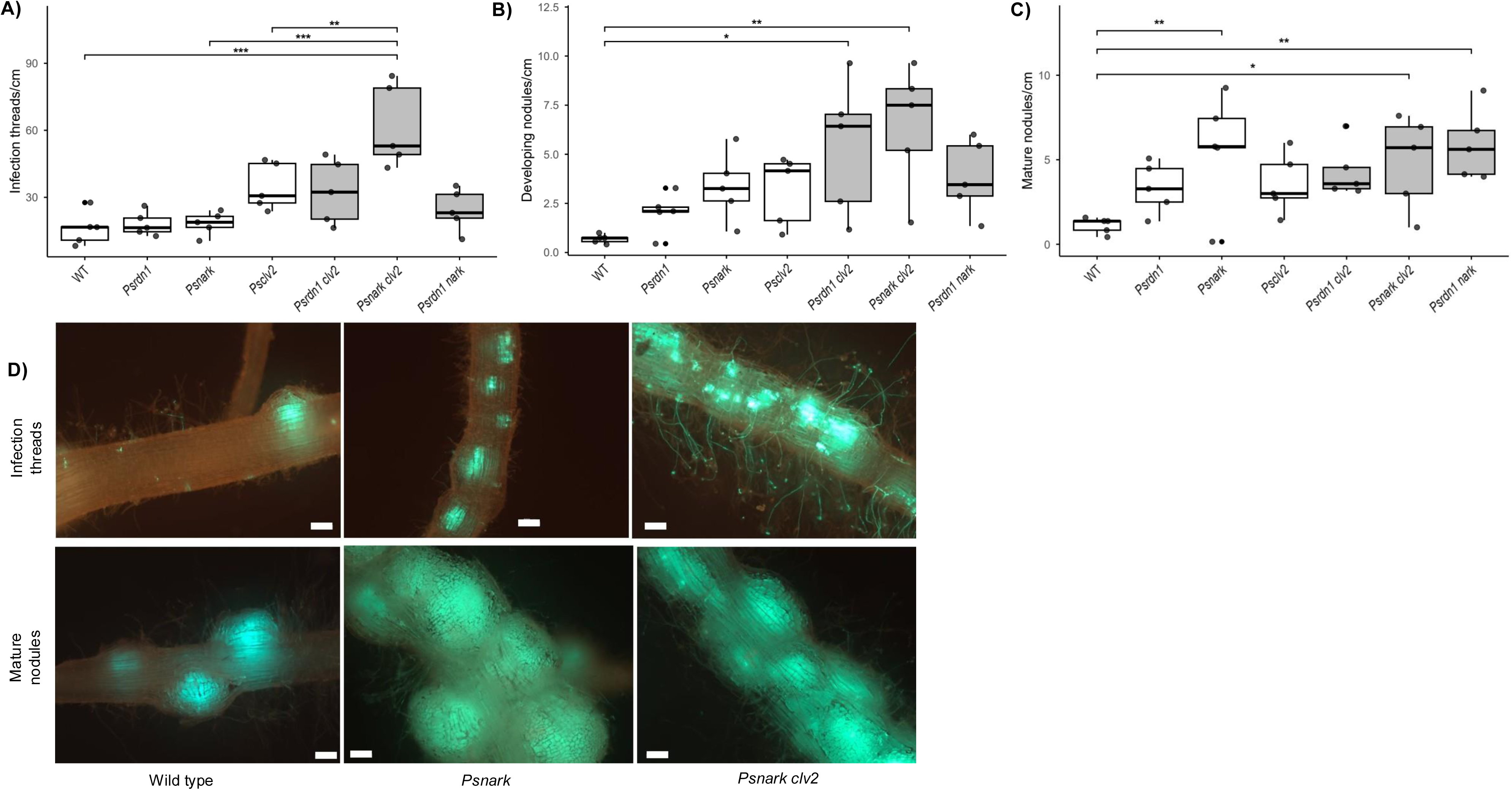
Nodulation phenotypes of wild type (WT; Frisson), *Psrdn1*, *Psnark, Psclv2, Psrdn1 clv2*, *Psnark clv2* and *Psrdn1 nark* mutant peas. (A) Number of infection threads per cm of root. (B) Number of developing nodules per cm of root. (C) number of mature nodules per cm of root. (D) Photograph of individual secondary roots showing infection threads connected to developing nodules and mature nodules taken with GFP filter, GFP indicates *Rhizobium leguminosarum*, scale bars = 200 µm. For (A, B, C) Boxes show interquartile range, with the inner line representing the median of each treatment. Outliers fall beyond the whiskers, which encompasses up to 1.5 times the length of the interquartile range. (A-C) Statistical analysis performed using one-way linear model. Post hoc analyses compared mutant lines to the control (WT) using Dunnett’s test via estimated marginal means (EMM). Assumptions were tested by Shapiro–Wilk and Levene’s test. Significance between these genotypes is shown as *, **, *** for p <0.05, 0.01, 0.001, respectively. Additionally, each double mutant was compared to its component single mutants with Holm adjustment for testing within families.

Due to relatively high variation, no single mutants displayed a significant increase in number of developing nodules compared to wild type plants, although the trend was for single mutants to have more developing nodules (Fig 5B). *Psrdn1 clv2* and *Psnark clv2* mutants did have significantly more developing nodules than wild type, but were not significantly different from single mutant parents (Fig 5B). All three single mutant plants formed significantly more mature nodules than wild type. However, *Psnark* single mutant plants formed strikingly more mature nodules than other single mutants; 6-fold more mature nodules than wild type (Fig 5C). Fluorescent imaging revealed that the mature root of *Psnark* was completely covered in nodules and lacked space for additional nodules to emerge (Fig 5D). *Psnark clv2* and *Psrdn1 nark* double mutant plants also formed an excessive number of mature nodules, not significantly different from *Psnark*. Given the almost complete saturation of the *Psnark* root with mature nodules, it is unlikely that any additive effect of additional mutations could be observed on a *Psnark* mutant background. Thus, it is difficult to conclude if *Ps*NARK and *Ps*CLV2 act in same or partially independent pathways to suppress mature nodule number. However, it is clear that *Ps*CLV2 and *Ps*RDN1 likely act in the same pathway to suppress nodule number, as no additive effect on mature nodule number was observed in *Psclv2 rdn1* compared to single mutant *Psclv2* and *Psrdn1* single mutant plants (Fig 5C).

Another way to examine the relationship between *Ps*NARK and *Ps*CLV2 is by grafting between *Psnark*, *Psclv2* and wild type lines. As previous grafting studies have revealed that both *Ps*NARK and *Ps*CLV2 receptors act in the shoot (Sagan and Duc, 1996), grafting between the mutants can highlight whether these receptors are likely to act together and/or in parallel pathways in the shoot to suppress nodulation in the root. Grafting between receptor mutants of the AON pathway has not been previously reported in any species. Grafting a wild type scion to a *Psclv2* rootstock suppressed nodulation in the *Psclv2* rootstock to the same level as seen in a wild type self-grafted plant (Fig 6B). However, a *Psnark* scion was unable to suppress nodulation in a *Psclv2* rootstock and actually led to higher nodule number than was observed in *Psclv2* self-grafted plant (Fig 6B). This suggests that in the shoot, *Ps*CLV2 cannot act independently of *Ps*NARK to suppress nodulation.

**Figure 6.**
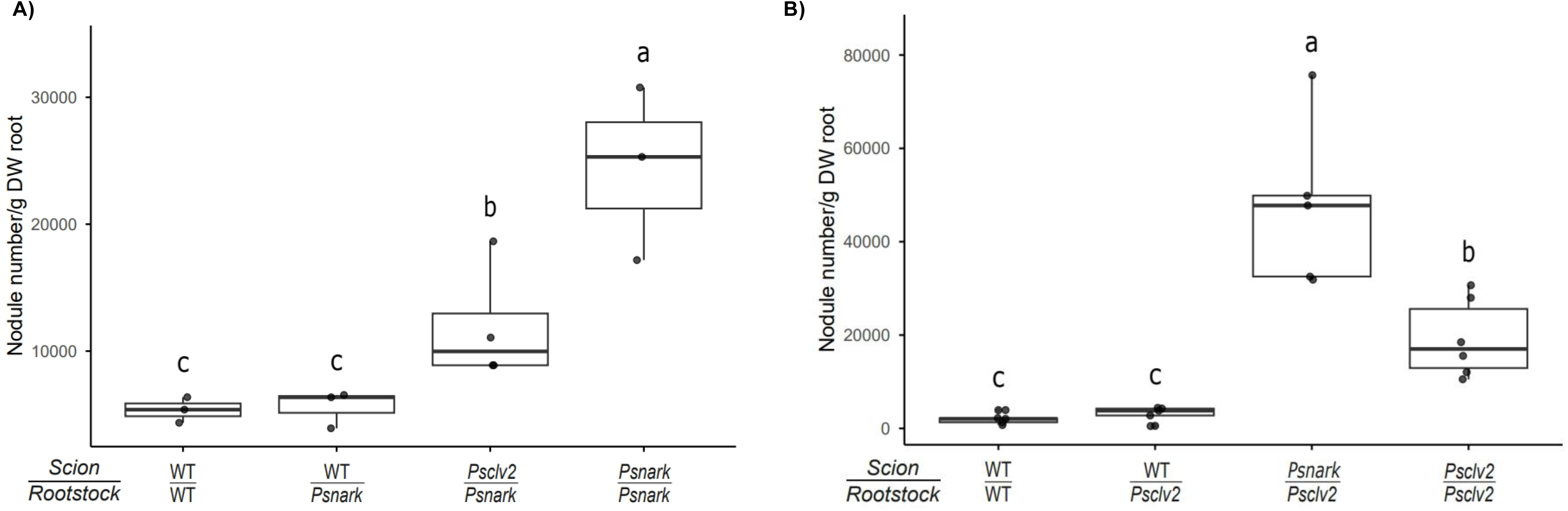
Nodulation in reciprocal grafts between wild type (WT; Frisson), *Psnark* and *Psclv2* mutant peas. (A, B) Number of mature nodules per g root, scion and rootstock are indicated. Boxes show interquartile range, with the inner line representing the median of each treatment. Outliers fall beyond the whiskers, which encompasses up to 1.5 times the length of the interquartile range. Letters above each box represent significance values generated by one-way ANOVA comparing treatments within panel. (n= 4-5).

We also examined the influence of wild type or *Psclv2* scion on *Psnark* rootstock. Grafting a wild type scion to a *Psnark* rootstock suppressed nodulation in the *Psnark* rootstock such that it resembled a wild type self-grafted plant (Fig 6A). Grafting to a *Psclv2* scion caused reduction in nodule number in *Psnark* rootstock, resulting in nodule number that was intermediate between the effect of grafting to a wild type scion and a *Psnark* scion (Fig 6A). This suggests that *Ps*NARK may suppress nodulation in part by acting with *Ps*CLV2, but also by acting independently of *Ps*CLV2.

## Discussion

This study has established a clear model for the key players in the AON system of pea (Fig 7) that shares many similarities, but also some key differences, with other model legumes. Two Rh-CLE peptides, *Ps*CLE12 and *Ps*CLE13 (Fig 1), that are likely orthologs of *Mt*CLE12 and *Mt*CLE13 respectively, systemically suppress nodule number in pea (Figs 2-3). The systemic suppression of nodule number requires both CLV1 ortholog *Ps*NARK and the LRR receptor-like protein *Ps*CLV2 (Figs 2, 3). Interestingly, both peptides require *Ps*RDN1 to suppress nodule number (Fig 4), indicating that both likely require arabinosylation for their activity.

**Figure 7.**
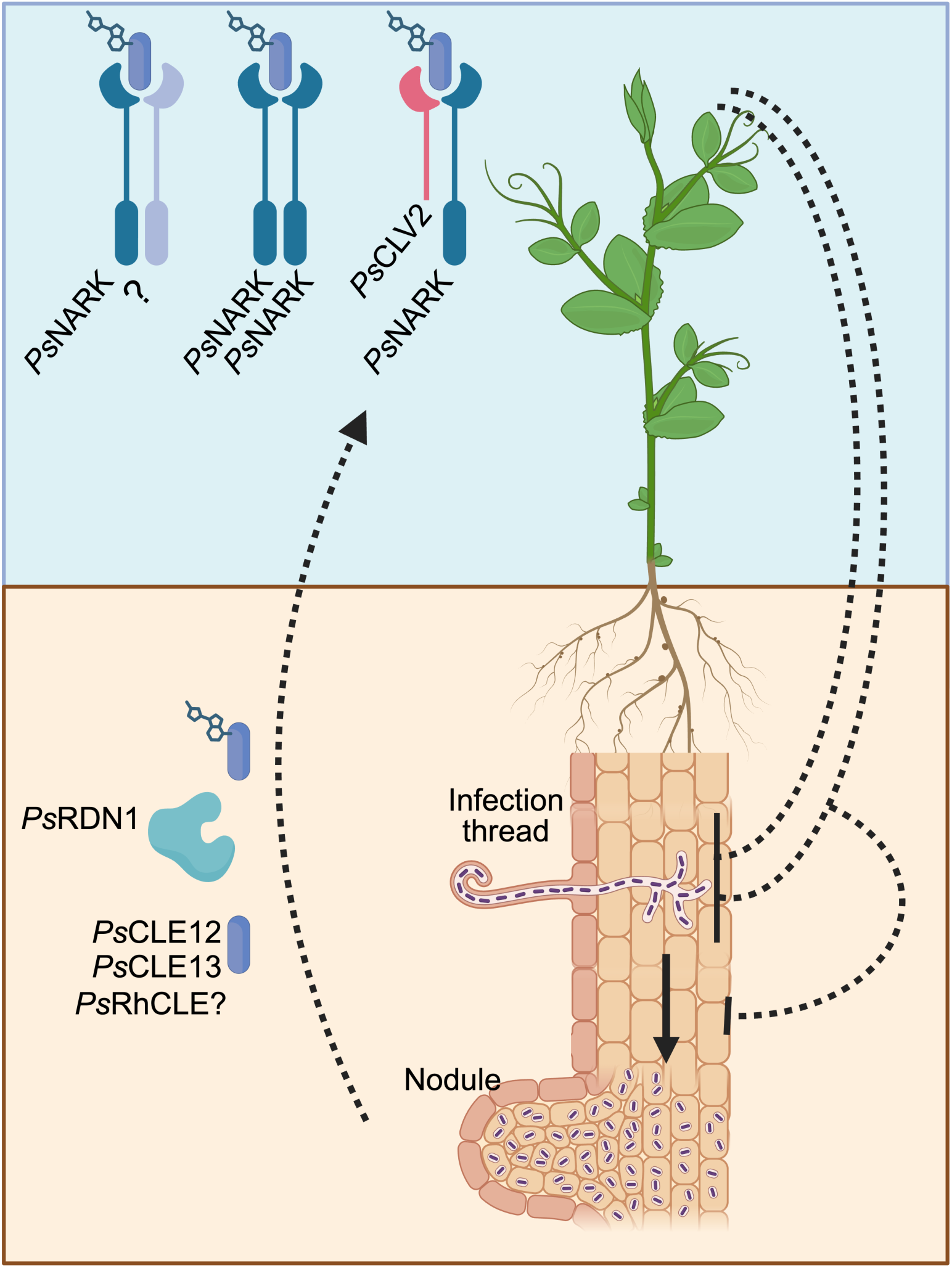
Model of the proposed CLE peptide signalling pathway regulating nodulation in pea. Rhizobial-induced Rh-CLEs *Ps*CLE12 and *Ps*CLE13 are arabinosylated via *Ps*RDN1 and move to shoot. Other as yet undiscovered RhCLE may also act in this pathway. In the shoot *Ps*CLE12 and *Ps*CLE13 are perceived by *Ps*NARK, which can act by complexing with *Ps*CLV2 but also by complexing with itself or other unknown shoot acting receptors. Shoot-to-root signal(s) then suppress both infection and nodule organogenesis in the root. (Figure produced with BioRender).

This is in contrast to *Medicago*, in which both *Mt*CLE12 and *Mt*CLE13 require arabinosylation for activity but only *Mt*CLE12 requires *Mt*RDN1 (Imin *et al*., 2018; Kassaw *et al*., 2017). The lack of additive nodulation phenotype in *Psclv2 rdn1* double mutant plants compared to single mutant parents (Fig 5) indicates that *Ps*RDN1 acts in the same pathway as *Ps*CLV2, likely by arabinosylating CLE peptides that are then perceived by CLV2. This is the first direct evidence that RDN1 and CLV2 act in the same pathway to control nodulation and contrasts with their role in parallel pathways in control of mycorrhizal colonisation in tomato (Wulf *et al*., 2023).

We examined how the AON system influences the nodulation process; infection in the epidermis and nodule organogenesis in the inner cortex. We found that disruption of *Ps*NARK and *Ps*CLV2 led to mild but in some cases significant increases in infection thread and nodule number (Fig 2, 3). However, it is important to point out that although nodule number was always elevated in these mutants, infection thread number was only elevated in some experiments. Furthermore, we found that overexpression of *PsCLE12* or *PsCLE13* only suppressed developing and/or mature nodule number and had no significant effect on infection thread number across several experiments (Fig 2-4). This is in contrast to other studies that have examined the impact of CLE overexpression on rhizobial infection, which found that in *Medicago MtCLE35* overexpression suppressed infection thread number (Moreau *et al*., 2021) and in *Lotus* overexpression of *LjCLE-RS1* or *LjCLE-RS2* suppressed the hyper-infection thread phenotype of *daphne/nin* mutant (Yoro *et al*., 2019b). Thus, our results suggest CLE peptides in addition to *Ps*CLE12 and *Ps*CLE13 may signal through *Ps*NARK and *Ps*CLV2 to suppress infection thread formation. Together these results suggests that the AON system in pea may influence both the formation of infection threads and the concomitant induction of nodule organs associated with some, but not all, infection threads (Fig 7), consistent with models in *Medicago* and *Lotus* (Imin *et al*., 2018; Moreau *et al*., 2021; Yoro *et al*., 2019b).

Studies in other model legumes have indicated downstream transcriptional targets of the AON system in roots are TML1 and TML2. In the days following exposure to rhizobia, we found that enhanced *PsCLE12* and *PsCLE13* expression did not precede upregulation of *PsTML1* and *PsTML2* (Fig 1). Indeed, unlike upregulation of *MtTML1* and *MtTML2* expression in lines overexpressing *MtCLE12* and *MtCLE13* in *Medicago* (Gautrat *et al*., 2019), we found over expression of *PsCLE12* and *PsCLE13* systemically suppressed nodulation but did not lead to elevated *PsTML1* or *PsTML2* expression (Suppl Fig 1,2). This suggests that in pea additional Rh-CLE peptides or other mechanisms may regulate *PsTML1/2* expression. As has been previously reported for *Medicago* in an RNAseq timecourse (Schnabel *et al*., 2023), we found *PsTML1* and *PsTML2* expression was not altered in *Psrdn1* mutants and this was consistent with no change in expression profile in *Psclv2* (Suppl Fig 1, 2). However, a small decrease in *PsTML1* and *PsTML2* expression was observed in *Psnark* mutants (Suppl Fig 1, 2). Taken together, lack of a consistent influence of all AON elements examined (CLE peptides, receptor and HPAT modifying enzymes) on *PsTML1* and *PsTML2* expression across many independent experiments suggests that players in addition to TML1 and TML2 integrate AON signalling in pea.

To examine the interaction between *Ps*NARK and *Ps*CLV2, the putative CLE receptors in pea, we performed double mutant studies and grafting between single mutants. The additive infection thread phenotype of *Psnark clv2* mutants compared to single mutant parents (Fig 5a, d), indicates these receptors are likely to act at least partially independently. The fact that nodule number was not additive may be due to the observation that almost all available root space for nodules was already occupied with nodules in any plant with the *Psnark* mutation (Fig 5 B, C, E). In *Lotus*, the same double mutant *har clv2* has also been reported to display no additive effect on nodule number, although as no quantitative data on nodules or infection threads were reported it is difficult to directly compare experiments across species. Grafting between single mutant *Psnark* and *Psclv2* mutants offers an elegant way to examine the potential interaction of these receptors. We found that *Psclv2* shoots could partially rescue the nodule number of *Psnark* roots, but not vice versa, indicating *Ps*NARK can suppress nodulation in part by acting with *Ps*CLV2 but also by acting independently of *Ps*CLV2. This means *Ps*NARK may act as a homo-dimer and/or heterodimer with an unknown co-receptor, while *Ps*CLV2 likely acts as a heterodimer with *Ps*NARK (Fig 7). Potential co-receptors identified in other legume species outlined in the introduction include KLAVIER, CORYNE and BAM2, although no mutants in these genes have been identified in pea. As outlined above, *Ps*RDN1 is likely to act in same pathway as *Ps*CLV2, and the lack of additive phenotype in infection or nodule number in the *Psnark rdn1* double mutant (Fig 5) suggests that *Ps*RDN1 also acts upstream of *Ps*NARK. Together with the fact that all three elements are required for *Ps*CLE12 and *Ps*CLE13 action, our current model of AON in pea involves several parallel pathways suppressing infection and nodule organogenesis (Fig 7).

The intriguing observation that *Psnark* mutant shoots were unable to completely restore nodule suppression in *Psclv2* roots, as occurs in *Psclv2* roots grafted to wild type scion (Fig 6 a), suggests that active compensation may be occurring in the signalling components of the AON system of pea. Active compensation, or genetic buffering, is a common feature of the CLE peptide signalling system that controls shoot apical meristem development, a pathway that shares common genes with the AON/AOM system (reviewed by Scott *et al*., 2024).

Indeed, in pea, examination of the shoot phenotypes of *Psnark clv2* double mutants revealed loss of *PsNARK* likely leads to active compensation by another shoot signalling element (not *Ps*CLV2) to promote shoot apical meristem proliferation (Scott *et al*., 2024). Recently, a study in *Medicago* suggested that BAM2, a putative CLE receptor, may compensate for the lack of CLV1-ortholog *Mt*SUNN in the roots to suppress nodulation (Thomas and Frugoli, 2023). Therefore, mutation of *Ps*NARK in the shoot may lead to altered expression/activity of other shoot acting AON elements (such as other CLE receptors) that regulate nodulation. Future studies could use tissue-specific transcriptomics combined with transgenic studies to identify potential candidates in this potential AON genetic buffering.

CLE peptide signalling is an emerging area of interest for understanding the control of many complex processes, including the control of symbioses. The emerging consensus is that in symbioses CLE signalling does not operate via simple root-shoot-root loops, but are likely made up of many parallel signalling loops with key overlaps (for review see Chaulagain and Frugoli, 2021; Roy and Muller, 2022; Scott *et al*., 2024) and even symbiont derived peptides (Bashyal *et al*., 2025). Future work should aim to uncover other key players in AON and AOM, including additional root-to-shoot CLE peptides, modifying enzymes, shoot receptors, shoot-to-root signals, and key targets including root integrating modules and downstream transcriptional targets. This understanding will underpin future optimisation of symbioses in crops.

## Supplementary data

**Supplemental Table 1-** Primer information for real time PCR.

**Supplemental Table 2-** DEGs in symbioses pathway in pea roots 10 days post inoculation (dpi) with rhizobia (vs 0 dpi). Colours indicate different stages of nodulation.

**Supplemental Figure 1.** Expression profiles of key autoregulation genes in wild type (WT; Frisson) and *Psnar*k, *Psclv2* or *Psrdn1* mutants transformed via hairy root transformation with control (*35S::GUS*) or *35S::PsCLE12* construct. Significance determined using Welch t-test or, when Shapiro or Levene assumptions not met, Wilcoxon rank-sum.

**Supplemental Figure 2.** Expression profiles of key autoregulation genes in wild type (WT; Frisson) and *Psnar*k, *Psclv2* or *Psrdn1* mutants transformed via hairy root transformation with control (*35S::GUS*) or *35S::PsCLE13* construct. Significance determined using Welch t-test or, when Shapiro or Levene assumptions not met, Wilcoxon rank-sum.

## Acknowledgments

We gratefully acknowledge technical assistance of Fu Rong Mah and Alannah Mannix and expert plant husbandry of Sarah Kane and Tracey Winterbottom.

## Author Contribution Statement

EF conceived and designed the project. TS, KW, AC-L, EF and KV conducted the experiments. TS, EF, AC-L, KW and KV analysed the data. EF wrote the manuscript with input from TS, KV and JBR. All authors read and approved the manuscript.

## Data availability

The RNA-seq datasets generated for this study have been deposited in the NCBI Sequence Read Archive (SRA) under accession number PRJNA1327600.

