## Supplementary material for "Parallel CLE peptide signaling pathways control nodulation in pea": Table S1

Supplemental Table 1. **List of primers used for qPCR.**

| Gene name | Accession | Primer |
|---|---|---|
| PsCLE12 | PsCam040153 | F - 3’- AGACTTGAAGGTGGTGGAAGAC-5’ |
| | | R - 3’- AAAGCCTATCTTCACCTGTAGGT-5’ |
| PsCLE13 | PsCam040702 | F - 3’-GGGTCGGTATACAAATCAAGTGC-5’ |
| | | R - 3’-GGTCCACCTGGTGAGAGTCT-5’ |
| PsTML1 | PsCam036214 | F1-5’- GGCGGCTTAGAAACAAACAC-3’ |
| | | R1-5’- TCTCATCCTGTCCACCAATAAC -3’ |
| PsTML2 | PsCam037480 | F4-5’- CACACATTTGTTTGTTTCAGGTATG-3’ |
| | | R4-5’- GATGAAGCTGATGCAAAGACAC-3’ |
| PsNIN | PsCam045001 | F4 – 5’-AAAGAGCATCGGTGTATGTCC -3’ |
| | | R4 – 5’- CCTCAGCACCTTGAACAGAA -3’ |
