## Supplementary figures and images for "Parallel CLE peptide signaling pathways control nodulation in pea"

### Fig S1

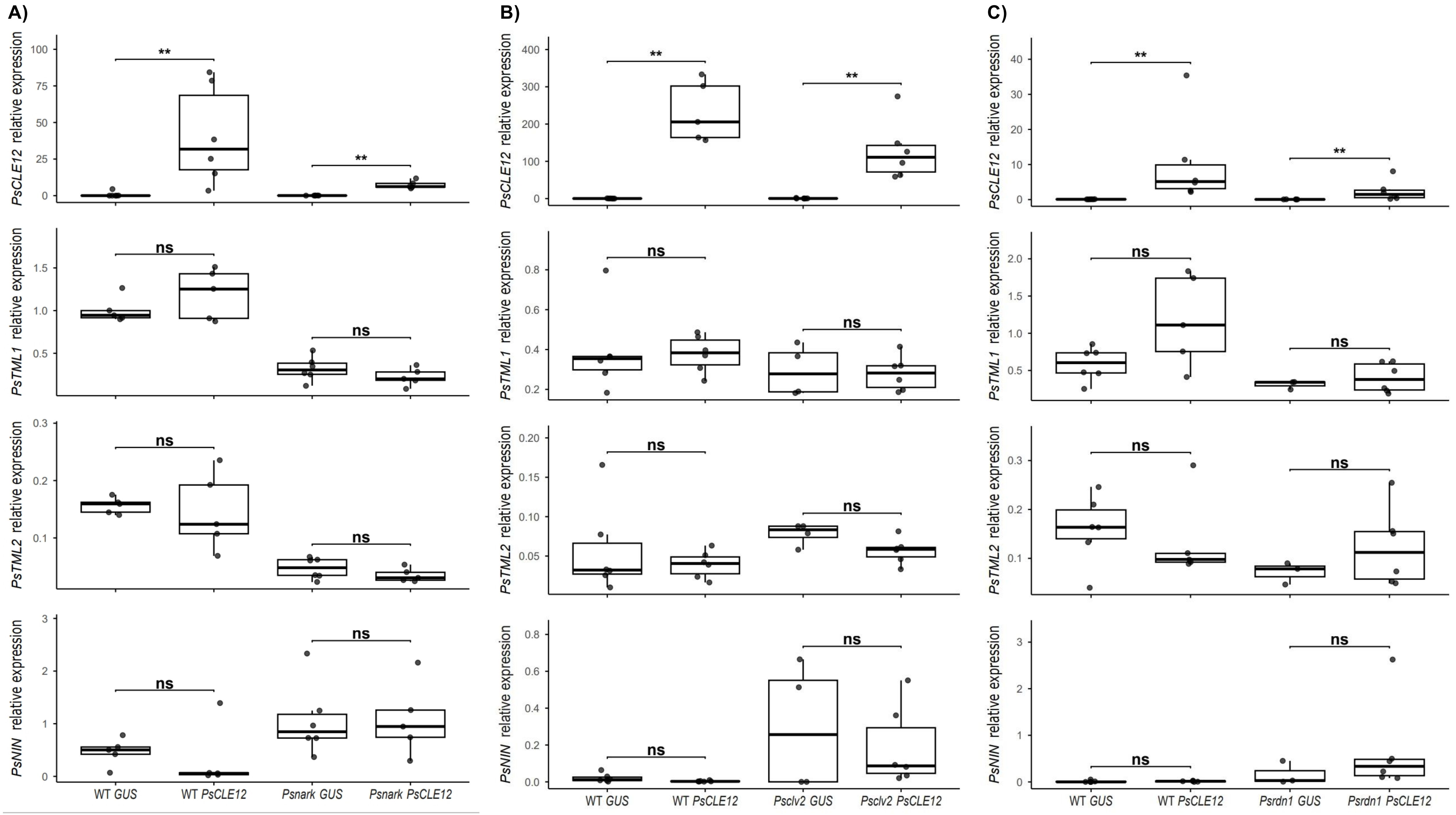

### Fig S2

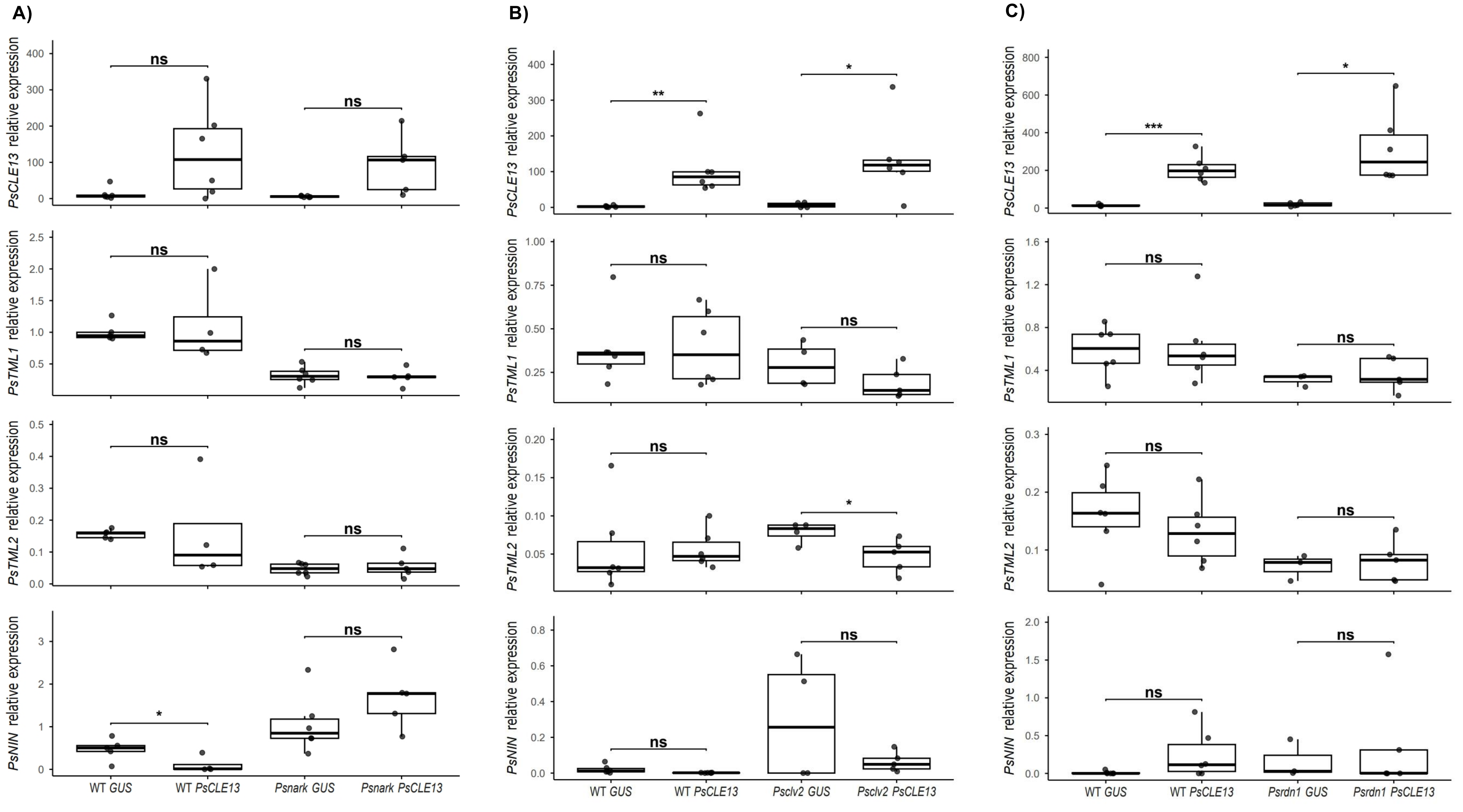
