## Supplementary material for "Parallel CLE peptide signaling pathways control nodulation in pea": Table S2

| Nodulation Stage | Gene name | Transcript_ID | PsatID | log2Fold Change | | q value |
|---|---|---|---|---|---|---|
| Early Signaling | NUCLEOPORIN 85 (NUP85) | PsCam000629 | Psat2g005000 | 0.178 | ↑ | 9.51E-04 |
| Early Signaling | DELLA 2 | PsCam016747 | Psat7g082120 | 0.267 | ↑ | 7.18E-04 |
| Early Signaling | SYMBIOTIC RECEPTOR KINASE (SYMRK),DOES NOT MAKE Infections 2 (DMI2) | PsCam031160 | Psat2g134120 | 0.375 | ↑ | 8.15E-12 |
| Early Signaling | DOES NOT MAKE Infections 1 (DMI1) | PsCam049032 | Psat1g222280 | 0.382 | ↑ | 5.86E-08 |
| Early Signaling | NFR5-interacting cytoplasmic kinase 4 (NiCK4) | PsCam000863 | Psat4g135400 | 0.446 | ↑ | 5.02E-07 |
| Early Signaling | SYMRK INTERACTING PROTEIN 1 (SIP1) | PsCam050140 | Psat5g008600 | 0.446 | ↑ | 1.66E-06 |
| Early Signaling | NOD FACTOR RECEPTOR (NFR5) | PsCam036765 | Psat2g162600 | 0.451 | ↑ | 1.53E-02 |
| Early Signaling | DELLA 3 | PsCam026903 | Psat1g025560 | 0.877 | ↑ | 1.05E-08 |
| Early Signaling | NODULATION SIGNALING PATHWAY 1 (NSP1) | PsCam035737 | Psat0s741g0280 | 0.898 | ↑ | 3.48E-15 |
| Early Signaling | CHITINASE 5 (CHIT5) | PsCam037392 | Psat0s5923g0040 | 1.086 | ↑ | 2.91E-20 |
| Early Signaling | INTERACTING PROTEIN DMI 3 | PsCam020896 | Psat2g142200 | 1.381 | ↑ | 2.67E-81 |
| Early Signaling | 3-HYDROXY-3-METHYLGLUTARYL COENZYME A REDUCTASE 1 (HMGR1) | PsCam020782 | Psat2g146800 | 1.628 | ↑ | 1.39E-24 |
| Early Signaling | PLANT U BOX PROTEIN 1 (PUB1) | PsCam033292 | Psat2g030880 | 2.949 | ↑ | 6.61E-108 |
| Early Signaling | EXOPOLYSACCHARIDE RECEPTOR (EPR3) | PsCam036987 | Psat2g094040 | 4.179 | ↑ | 8.04E-120 |
| Early Signaling | NOD FACTOR HYDROLASE 1 (NFH1) | PsCam038449 | Psat4g033880 | 7.485 | ↑ | 5.10E-127 |
| Early Signaling | SYMRK INTERACTING PROTEIN 2 (SIP2) | PsCam034787 | Psat7g263320 | -0.099 | ↓ | 2.00E-02 |
| Early Signaling | CHALCONE REDUCTASE (CHR) | PsCam033579 | Psat2g003880 | -0.880 | ↓ | 1.80E-76 |
| Early Signaling | BRUSH2 | PsCam009542 | Psat0s2056g0040 | -1.310 | ↓ | 7.95E-03 |
| Rizobial Infection | PHOSPHATIDYLINOSITOL 3-KINASE (PI3K) | PsCam059963 | Psat2g104920 | 0.161 | ↑ | 1.94E-02 |
| Rizobial Infection | TARGET OF RAPAMYCIN (TOR) | PsCam023168 | Psat4g228960 | 0.185 | ↑ | 3.68E-03 |
| Rizobial Infection | nodule specific REALLY INTERESTING NEW GENE (nsRING) | PsCam023219 | Psat7g010880 | 0.544 | ↑ | 2.26E-11 |
| Rizobial Infection | EXOCYST70H4 | PsCam036091 | Psat7g220480 | 0.619 | ↑ | 3.18E-17 |
| Rizobial Infection | NODULIN 5 (N5) | PsCam048664 | Psat2g010560 | 0.698 | ↑ | 2.06E-03 |
| Rizobial Infection | FLOTILLIN 4 (FLOT4)2 | PsCam012878 | Psat5g030840 | 1.365 | ↑ | 4.53E-26 |
| Rizobial Infection | FLOTILLIN 4 (FLOT4)1 | PsCam057726 | Psat5g030720 | 1.564 | ↑ | 1.66E-13 |
| Rizobial Infection | Rhizobial Infection Receptor-like Kinase1 | PsCam034400 | Psat5g113400 | 2.766 | ↑ | 5.78E-122 |
| Rizobial Infection | FLOTILLIN 4 (FLOT4)3 | PsCam057775 | Psat5g030760 | 3.253 | ↑ | 4.51E-03 |
| Rizobial Infection | VAPYRIN (VPY) | PsCam030841 | Psat1g090680 | 3.337 | ↑ | 1.66E-173 |
| Rizobial Infection | RHIZOBIUM-DIRECTED POLAR GROWTH (RPG) | PsCam035887 | Psat6g173600 | 4.686 | ↑ | 2.78E-48 |
| Rizobial Infection | CYSTATHIONE BETASYNTHASE 1 (CBS1) | PsCam038043 | Psat3g167920 | 7.744 | ↑ | 1.04E-12 |
| Rizobial Infection | SYMBIOTIC PROTEIN KINASE 1 (SPK1) | PsCam038220 | Psat0s4890g0040 | 8.713 | ↑ | 5.85E-43 |
| Rizobial Infection | NUCLEAR FACTOR YA1 | PsCam037673 | Psat6g092920 | 9.382 | ↑ | 3.99E-61 |
| Rizobial Infection | NODULE PECTATE LYASE (NPL) | PsCam034822 | Psat5g092760 | 12.521 | ↑ | 1.24E-24 |
| Rizobial Infection | SYMBIOTIC REMORIN1 (SymREM1) | PsCam031115 | Psat7g034920 | 13.862 | ↑ | 1.86E-30 |
| Rizobial Infection | Rboh(respiratory burst oxidase homologues, NADPH oxidase) (RbohB) | PsCam045017 | Psat3g011120 | -0.148 | ↓ | 1.50E-02 |
| Rizobial Infection | Rboh(NADPH-oxidase) (RbohA,RbohB) | PsCam055889 | Psat5g026680 | -0.184 | ↓ | 3.41E-02 |
| Rizobial Infection | 121F-specific p53 INDUCIBLE RNA (PIR) | PsCam049039 | Psat0s3583g0040 | -0.202 | ↓ | 2.37E-04 |
| Rizobial Infection | Nck-ASSOCIATED LIKE PROTEIN 1 (NAP1) | PsCam044865 | Psat7g129680 | -0.239 | ↓ | 1.80E-05 |
| Rizobial Infection | CALCIUM DEPENDANT PROTEIN KINASE 1 (CDPK1) | PsCam049094 | Psat2g154520 | -0.391 | ↓ | 1.45E-09 |
| Nodule Organogenesis | INCREASING NODULE SIZE 1 (INS1) | PsCam048191 | Psat6g051440 | 0.190 | ↑ | 6.18E-03 |
| Nodule Organogenesis | Soybean heterotrimeric G PROTEINS (Ga) | PsCam026885 | Psat6g021680 | 0.262 | ↑ | 1.28E-02 |
| Nodule Organogenesis | Digalactosyldiacylglycerol (DGDG) synthase 1 | PsCam048970 | Psat3g028880 | 0.265 | ↑ | 9.51E-05 |
| Nodule Organogenesis | BYPASS1 (BPS1)1 | PsCam034003 | Psat3g108800 | 0.285 | ↑ | 2.24E-06 |
| Nodule Organogenesis | ASSOCIATED MOLECULE WITH THE SH3 DOMAIN OF STAM (AMSH) | PsCam037176 | Psat1g036960 | 0.286 | ↑ | 9.77E-04 |
| Nodule Organogenesis | CELL CYCLE SWITCH GENE encoding a 52 kDa protein (CCS 52a) | PsCam055886 | Psat0s3705g0040 | 0.332 | ↑ | 1.56E-08 |
| Nodule Organogenesis | RESPONSE REGULATOR (RR5,9,11) | PsCam039920 | Psat5g220920 | 0.430 | ↑ | 3.55E-03 |
| Nodule Organogenesis | PLETHORA 1 (PLT1,PLT2,PLT3,PLT4)2 | PsCam037251 | Psat5g287880 | 0.432 | ↑ | 4.01E-02 |
| Nodule Organogenesis | CYTOKININ RESPONSE 1 (CRE1) | PsCam044895 | Psat7g004720 | 0.453 | ↑ | 5.13E-10 |
| Nodule Organogenesis | PLETHORA 1 (PLT1,PLT2,PLT3,PLT4)5 | PsCam036332 | Psat3g103160 | 0.473 | ↑ | 5.38E-03 |
| Nodule Organogenesis | BYPASS1 (BPS1)2 | PsCam042733 | Psat3g108760 | 0.530 | ↑ | 7.14E-07 |
| Nodule Organogenesis | BASIC HELIXâ€"LOOPâ€"HELIX 1 (BHLH1) | PsCam038780 | Psat5g050120 | 0.584 | ↑ | 2.07E-04 |
| Nodule Organogenesis | NUCLEAR FACTOR YC1 | PsCam053713 | Psat3g010480 | 0.689 | ↑ | 8.34E-32 |
| Nodule Organogenesis | NODULE ROOT (NOOT) | PsCam036654 | Psat3g077680 | 0.728 | ↑ | 7.70E-07 |
| Nodule Organogenesis | ISOFLAVONE SYNTHASE (IFS) | PsCam025494 | Psat7g118080 | 0.740 | ↑ | 1.85E-07 |
| Nodule Organogenesis | KNOTTED-1 LIKE TAKE HOMEDOMAIN proteins 4 | PsCam050692 | Psat0s149g0040 | 0.879 | ↑ | 2.13E-77 |
| Nodule Organogenesis | ANNEXIN1 (ANN1)2 | PsCam043064 | Psat4g147120 | 0.902 | ↑ | 5.07E-18 |
| Nodule Organogenesis | ANNEXIN1 (ANN1)1 | PsCam009798 | Psat4g191080 | 0.913 | ↑ | 2.39E-14 |
| Nodule Organogenesis | LUMPY InfectionS (LIN) | PsCam030891 | Psat0s1505g0120 | 1.033 | ↑ | 1.48E-65 |
| Nodule Organogenesis | PLETHORA 1 (PLT1,PLT2,PLT3,PLT4)4 | PsCam036983 | Psat2g089880 | 1.118 | ↑ | 3.93E-16 |
| Nodule Organogenesis | CYTOKININ OXIDASE,DEHYDROGENASE 3 (CKX3) | PsCam000181 | Psat4g015400 | 1.284 | ↑ | 5.22E-05 |
| Nodule Organogenesis | KNOTTED-1 LIKE TAKE HOMEDOMAIN proteins 2 | PsCam054829 | Psat5g030920 | 1.367 | ↑ | 2.60E-23 |
| Nodule Organogenesis | NIN LIKE PROTEIN (NLP1) | PsCam035627 | Psat0s498g0160 | 1.492 | ↑ | 7.62E-86 |
| Nodule Organogenesis | KNOTTED-1 LIKE TAKE HOMEDOMAIN proteins 1 | PsCam042941 | Psat6g028400 | 1.686 | ↑ | 4.37E-53 |
| Nodule Organogenesis | KNOTTED-1 LIKE TAKE HOMEDOMAIN proteins 3 | PsCam050922 | Psat4g033160 | 2.467 | ↑ | 2.66E-139 |
| Nodule Organogenesis | ETHYLENE RESPONSE FACTOR REQUIRED FOR NODULE DIFFERENTIATION (EFD) | PsCam044465 | Psat7g259160 | 3.034 | ↑ | 1.33E-83 |
| Nodule Organogenesis | PIN-FORMED(PIN2,PIN3,PIN4)2 | PsCam053961 | Psat0s1191g0040 | 3.707 | ↑ | 4.26E-02 |
| Nodule Organogenesis | NODULE INCEPTION (NIN) | PsCam045001 | Psat2g001120 | 4.588 | ↑ | 1.62E-185 |
| Nodule Organogenesis | ?-EXPANSIN B2 (EXPB2) | PsCam039362 | Psat3g159240 | 6.297 | ↑ | 1.99E-69 |
| Nodule Organogenesis | RECEPTOR FOR ACTIVATED C KINASE (RACK1)2 | PsCam044430 | Psat7g003840 | -0.289 | ↓ | 5.32E-05 |
| Nodule Organogenesis | SUNERGOS 1 | PsCam029241 | Psat0ss6317g0080 | -0.293 | ↓ | 5.37E-04 |
| Nodule Organogenesis | ?-CAROTENE HYDROXYLASE (BCH1,BCH2) | PsCam001554 | Psat0ss1860g0160 | -0.304 | ↓ | 6.95E-03 |
| Nodule Organogenesis | CLATHARIN HEAVY CHAIN1 | PsCam010939 | Psat5g129720 | -0.313 | ↓ | 1.02E-25 |
| Nodule Organogenesis | CELL DIVISION CYCLE16 (CDC16) | PsCam044252 | Psat4g140160 | -0.358 | ↓ | 3.58E-07 |
| Nodule Organogenesis | Soybean heterotrimeric G PROTEINS2 | PsCam027543 | Psat0s2465g0080 | -0.373 | ↓ | 4.86E-05 |
| Nodule Organogenesis | NUMEROUS InfectionS AND POLYPHENOLICS,LATERAL ROOT-ORGAN DEFECTIVE (NIP,LATD) | PsCam006704 | Psat6g033120 | -0.419 | ↓ | 5.70E-05 |
| Nodule Organogenesis | ACYL CARRIER PROTEIN (ACP) | PsCam001965 | Psat3g103240 | -0.442 | ↓ | 1.48E-11 |
| Nodule Organogenesis | PIN-FORMED(PIN2,PIN3,PIN4)3 | PsCam010676 | Psat6g043160 | -0.484 | ↓ | 3.28E-05 |
| Nodule Organogenesis | PHYTOCHROME B (PHYB) | PsCam035479 | Psat1g161560 | -0.502 | ↓ | 2.29E-11 |
| Nodule Organogenesis | Soybean G-BOX FACTOR 14-3-3 (SGF14c,SGF14l) | PsCam001796 | Psat7g071680 | -0.566 | ↓ | 8.58E-33 |
| Nodule Organogenesis | TRICOT (TCO) | PsCam004740 | Psat3g026520 | -0.639 | ↓ | 5.85E-11 |
| Nodule Organogenesis | LIKE-AUX1 (LAX2) | PsCam034535 | Psat3g130520 | -0.647 | ↓ | 8.75E-31 |
| Nodule Organogenesis | BASIC LEUCINE ZIPPER FAMILY PROTEIN (BZF) | PsCam037032 | Psat5g115640 | -0.784 | ↓ | 1.51E-15 |
| Nodule Organogenesis | PIN-FORMED(PIN2,PIN3,PIN4)1 | PsCam036694 | Psat4g014160 | -0.791 | ↓ | 2.42E-08 |
| Nodule Organogenesis | PENETRATION3 - LIKE | PsCam035382 | Psat5g295880 | -0.849 | ↓ | 1.12E-17 |
| Nodule Organogenesis | ISOPENTENYL TRANSFERASE (IPT3) | PsCam038941 | Psat6g060960 | -1.050 | ↓ | 7.41E-15 |
| Nodule Organogenesis | AUXIN RESPONSE FACTOR 16a (ARF8a,ARF8b) | PsCam031281 | Psat2g042360 | -1.065 | ↓ | 1.73E-60 |
| Autorregulation of Nodule number | ROOT DETERMINED NODULATION 1 (RDN1) | PsCam048511 | Psat0s3806g0040 | 0.170 | ↑ | 2.63E-02 |
| Autorregulation of Nodule number | NITRATE UNRESPONSIVE SYMBIOSIS 1 (NRSYM1) | PsCam048071 | Psat6g197400 | 0.173 | ↑ | 1.25E-03 |
| Autorregulation of Nodule number | SUPERNUMERIC NODULE NUMBER (SUNN) | PsCam035758 | Psat7g183240 | 0.239 | ↑ | 1.13E-04 |
| Autorregulation of Nodule number | CORYNE (CRN) | PsCam009582 | Psat3g190240 | 0.439 | ↑ | 1.04E-07 |
| Autorregulation of Nodule number | COMPACT ROOT ARCHITECTURE 2 (CRA2) | PsCam042487 | Psat5g091280 | 0.651 | ↑ | 3.18E-14 |
| Autorregulation of Nodule number | **TOO MUCH LOVE (TML1,TML2)2** | **PsCam037480** | **Psat1g087960** | **0.926** | ↑ | **6.73E-17** |
| Autorregulation of Nodule number | **TOO MUCH LOVE (TML1,TML2)1** | **PsCam036214** | **Psat3g169800** | **0.972** | ↑ | **2.20E-62** |
| Autorregulation of Nodule number | CALCIUM DEPENDANT PROTEIN KINASE 3 (CPK3) | PsCam030792 | Psat5g178920 | -0.160 | ↓ | 5.09E-04 |
| Autorregulation of Nodule number | CLAVATA 2 (CLV2) | PsCam004573 | Psat3g104800 | -0.184 | ↓ | 2.32E-02 |
| Autorregulation of Nodule number | BES1,BZR1 HOMOLOG LIKE1 | PsCam023755 | Psat2g161520 | -0.220 | ↓ | 1.26E-04 |
| Autorregulation of Nodule number | KLAVIER (KLV)4 | PsCam043978 | Psat1g067160 | -0.265 | ↓ | 1.87E-04 |
| Autorregulation of Nodule number | NODULE NUMBER CONTROL 1 (NNC1) | PsCam042790 | Psat7g221800 | -0.492 | ↓ | 2.22E-06 |
| Bacterial Maturation | FERRITINS (FER2,FER3)2 | PsCam007305 | Psat7g247120 | 0.577 | ↑ | 7.47E-11 |
| Bacterial Maturation | FERRITINS (FER2,FER3)1 | PsCam058026 | Psat2g030280 | 1.037 | ↑ | 7.89E-65 |
| Bacterial Maturation | DEFECTIVE IN Nitrogen Fixation1 (DNF1) | PsCam051909 | Psat5g204840 | 1.354 | ↑ | 2.58E-46 |
| Bacterial Maturation | DEMETER (DME) | PsCam054805 | Psat6g177080 | 3.344 | ↑ | 8.71E-102 |
| Bacterial Maturation | DEFECTIVE IN Nitrogen Fixation2 (DNF2) | PsCam034802 | Psat7g125720 | 6.970 | ↑ | 8.13E-101 |
| Bacterial Maturation | STATIONARY ENDOSYMBIONT NODULE 1 (SEN1) | PsCam030926 | Psat7g097120 | 14.342 | ↑ | 3.01E-32 |
| Symbiosome Formation | AUTOPHAGY RELATED PROTEIN | PsCam050278 | Psat5g217520 | 0.313 | ↑ | 2.39E-07 |
| Symbiosome Formation | VESICLE ASSOCIATED MEMBRANE PROTEIN (VAMP721d,AMP721e) | PsCam001433 | Psat1g170080 | 0.422 | ↑ | 9.45E-11 |
| Symbiosome Formation | SYNAPTOGAMIN (SYT1,SYT2,SYT3)3 | PsCam029204 | Psat6g181920 | 0.589 | ↑ | 2.13E-10 |
| Symbiosome Formation | ACTIN RELATED PROTEIN (ARP2,ARP3) | PsCam049983 | Psat7g059080 | -0.149 | ↓ | 6.23E-03 |
| Symbiosome Formation | SYNTAXIN OF PLANTS 71 (SYP71) | PsCam051243 | Psat0s967g0040 | -0.289 | ↓ | 7.89E-10 |
| Symbiosome Formation | SYNAPTOGAMIN (SYT1,SYT2,SYT3)1 | PsCam049122 | Psat0s2115g0040 | -0.308 | ↓ | 2.35E-10 |
| Symbiosome Formation | Rab (Ras-related proteins in brain) GTPases1 | PsCam043528 | Psat7g250320 | -0.313 | ↓ | 7.03E-10 |
| Symbiosome Formation | Rab (Ras-related proteins in brain) GTPases2 | PsCam024934 | Psat7g141120 | -0.363 | ↓ | 6.04E-15 |
| Symbiosome Formation | SYNTAXIN OF PLANTS 132 (SYP132) | PsCam050501 | Psat5g264800 | -0.368 | ↓ | 5.04E-19 |
| Symbiosome Formation | Rab (Ras-related proteins in brain) GTPases Rabat | PsCam054893 | Psat7g211160 | -0.459 | ↓ | 3.18E-21 |
| Symbiosome Formation | SYNAPTOGAMIN (SYT1,SYT2,SYT3)2 | PsCam009630 | Psat6g003240 | -0.558 | ↓ | 7.65E-14 |
| Symbiosome Formation | TONOPLAST INTRINSIC PROTEIN 1g (TIP1g) | PsCam043552 | Psat7g21960 | -0.989 | ↓ | 1.20E-03 |
| Nodule Metabolism and Transport | SUCROSE SYNTHASE (SUCS1) | PsCam045132 | Psat4g019440 | 0.219 | ↑ | 9.52E-07 |
| Nodule Metabolism and Transport | NODULIN HOMEOBOX (NDX1,NDX2)2 | PsCam048982 | Psat5g282480 | 0.301 | ↑ | 7.55E-08 |
| Nodule Metabolism and Transport | PHOSPHATE TRANSPORTER 5 (PT5)2 | PsCam012944 | Psat3g057880 | 0.788 | ↑ | 1.97E-64 |
| Nodule Metabolism and Transport | METAL TOLERANCE PROTEIN 2 (MTP2) | PsCam000872 | Psat7g211200 | 0.891 | ↑ | 1.40E-19 |
| Nodule Metabolism and Transport | NRT1,PTR Family (NPF8.6) | PsCam004882 | Psat5g161760 | 3.128 | ↑ | 7.07E-04 |
| Nodule Metabolism and Transport | MOLYBDATE TRANSPORTER TYPE 1.2 | PsCam038189 | Psat6g031200 | 3.136 | ↑ | 1.03E-04 |
| Nodule Metabolism and Transport | SULPHATE TRANSPORTER 1 (SST1) | PsCam044086 | Psat1g040440 | 6.818 | ↑ | 1.43E-180 |
| Nodule Metabolism and Transport | NUCLEOLAR,MITOCHONDRIAL PROTEIN INVOLVED IN NODULATION (NMN) | PsCam031324 | Psat4g227240 | 6.873 | ↑ | 1.31E-183 |
| Nodule Metabolism and Transport | MULTIDRUG AND TOXIC COMPOUND EXTRUSION 1 (MATE1)3 | PsCam045540 | Psat4g191600 | 7.094 | ↑ | 2.82E-02 |
| Nodule Metabolism and Transport | MOLYBDATE TRANSPORTER TYPE 1.3 | PsCam033395 | Psat7g150560 | 7.588 | ↑ | 1.20E-135 |
| Nodule Metabolism and Transport | COPPER TRASPORTER 1 (COPT1) | PsCam044654 | Psat7g241040 | 8.127 | ↑ | 9.40E-09 |
| Nodule Metabolism and Transport | AMMONIUM TRANSPORTER 1.1 (AMT1.1) | PsCam000227 | Psat1g024160 | -0.108 | ↓ | 4.33E-02 |
| Nodule Metabolism and Transport | MULTIDRUG AND TOXIC COMPOUND EXTRUSION 1 (MATE1)1 | PsCam000259 | Psat5g027560 | -0.433 | ↓ | 1.54E-05 |
| Nodule Metabolism and Transport | MULTIDRUG AND TOXIC COMPOUND EXTRUSION 1 (MATE1)2 | PsCam038898 | Psat7g212600 | -0.535 | ↓ | 1.26E-08 |
| Nodule Metabolism and Transport | SUCROSE SYNTHASE (SUS1,SUS3) | PsCam044001 | Psat1g139760 | -0.644 | ↓ | 1.11E-146 |
| Nodule Metabolism and Transport | Natural Resistance-Associated Macrophage Protein1 (NRamp1) | PsCam012243 | Psat5g085960 | -1.357 | ↓ | 6.04E-09 |
| Nodule Metabolism and Transport | ZINC IRON PERMEASE 6 (ZIP6) | PsCam030944 | Psat7g132480 | -2.695 | ↓ | 8.30E-11 |
| Senescence | STAYGREEN (SGR) | PsCam001217 | Psat2g181040 | 0.222 | ↑ | 1.68E-02 |
| Senescence | CYSTEINE PROTEASE 15a (CYP15a) | PsCam014769 | Psat0s3831g0160 | 0.308 | ↑ | 3.79E-04 |
| Senescence | SERPIN (SER6)1 | PsCam038479 | Psat5g046600 | 0.417 | ↑ | 8.89E-03 |
| Senescence | VACUOLAR PROCESSING ENZYME (VPE) | PsCam004978 | Psat6g018800 | 0.589 | ↑ | 3.87E-09 |
| Senescence | CYSTEINE PROTEINASE 6 (CP6)2 | PsCam042142 | Psat2g153200 | 1.184 | ↑ | 2.07E-04 |
| Senescence | NAM,ATAF,CUC-encoding 969 (NAC969) | PsCam051494 | Psat7g159400 | 2.015 | ↑ | 2.68E-06 |
| Senescence | COLD ACCLIMATION SPECIFIC (CAS31) | PsCam007366 | Psat1g038720 | -4.548 | ↓ | 6.42E-04 |
| Defence | GLUTAMINE SYNTHETASE (GS),NADH?dependent GLUTAMATE SYNTHASE (NADH?GOGAT) | PsCam006801 | Psat6g037200 | 0.598 | ↑ | 3.82E-29 |
| Defence | STEAROYL-ACYL CARRIER PROTEIN DESATURASE ISOFORM C (SACPD-C) | PsCam043237 | Psat6g040040 | 1.791 | ↑ | 6.13E-22 |
| Defence | REGULATOR OF SYMBIOSOME DIFFERENTIATION (RSD) | PsCam031274 | Psat3g136520 | 11.756 | ↑ | 6.06E-07 |
| Defence | symbiotic CYSTEINE RICH KINASE (SymCRK) | PsCam030971 | Psat5g107080 | 13.568 | ↑ | 8.52E-24 |
| Defence | Glycine sojae 52 (GS52) | PsCam012963 | Psat3g090960 | -0.604 | ↓ | 2.32E-04 |
| Defence | ETHYLENE RESPONSE FACTOR 1 (ERF1) | PsCam038402 | Psat6g125480 | -1.077 | ↓ | 5.24E-15 |
